# *DIRC3* and close to *NABP1* Genetics Polymorphisms correlated with Prognostic Survival in Patients with Laryngeal Squamous Cell Carcinoma

**DOI:** 10.1101/042978

**Authors:** Zhen Shen, Wanli Ren, Yanxia Bai, Zhengshuai Chen, Jingjie Li, Bin Li, Tianbo Jin, Peilong Cao, Shao Yuan

**Affiliations:** Department of Otolaryngology&head neck, The First Affiliated Hospital of Xi'an Jiaotong University, Xi'an, Shaanxi 710061, China; Department of Pathology, The First Affiliated Hospital of Xi'an Jiaotong University, Xi'an, Shaanxi 710061, China; School of Life Sciences, Northwest University, Xi’an, Shaanxi 710069, China; National Engineering Research Center for Miniaturized Detection Systems, Xi’an 710069, China

**Keywords:** *DIRC3*, LSCC, *NABP1*, overall survival, prognostic factors

## Abstract

Laryngeal squamous cell carcinoma (LSCC) is one of the most common and aggressive malignancies in the upper digestive tract that has a high mortality rate and a poor prognosis. Prognostic factors were determined through multivariate Cox regression analysis. The overall survival rates were calculated by the Kaplan-Meier method. The SPSS statistical software package version 17.0 (SPSS Inc., Chicago, IL, USA) was used for all analyses. Median follow-up was 38 (range 3-122) months and the median survival time was 48 months. We adjusted to confounding factors (total laryngectomy, poor differentiation, T3-T4 stage, N1-N2 stage, III-IV TNM stage) into multivariate Cox proportional hazards model, we confirmed rs11903757 GT genotype (HR = 2.036; 95% CI, 1.071 - 3.872; p = 0.030) and rs966423 TT genotype (HR = 11.677; 95% CI, 3.901 - 34.950; p = 0.000) were significantly correlated with prognostic survival of patients with LSCC compared with rs11903757 TT genotype and rs966423 CC genotype, respectively. Our research provided new evidence for patients with LSCC, it seemed to be the first that demonstrated rs11903757 GT genotype on chromosome 2q32.3 close to *NABP1* and rs966423 TT genotype in the intron region of *DIRC3* on chromosome 2q35 predict poor prognostic survival in patients with LSCC.

## Introduction

Laryngeal squamous cell carcinoma (LSCC) is one of the most common and aggressive malignancies in the upper digestive tract that has a high mortality rate and a poor prognosis^1, 2^. Carcinomas of the upper aerodigestive tract represent a major problem in modern health care. The development of tumors is a complicated process which involves multiple genes and steps, and they are the result of the inter coordination and combined action of numerous factors. This is no exception of LSCC. Given the fundamental role the larynx plays in human speech and communication, determining the optimal management of laryngeal cancer is critical. For LSCC patients, the recurrence and metastasis of LSCC constitute the primary life-threatening reasons. This is primarily due to the lack of early diagnosis, leading to delayed implementation of suitable treatment and frequent recurrence. In China, the incidence of LSCC has been rising gradually, especially in the northeast part. The potentially high incidence of morbidity and incommensurably low cure rate, as expected, require searching for new diagnostic procedures and carcinogenic factors of LSCC ^3-5^. Despite significant advances in surgery and radio therapeutic techniques, as well as the attempt to use new chemotherapy drugs, the 5-year relative survival rates for patients with LSCC have not markedly improved. The mortality rate of LSCC is still high with 1.2 cases per 100,000 persons^6^. Although great progress has been achieved in the study on laryngeal cancer, there are no ideal biomarkers for the determination of prognosis and guidance of treatment in laryngeal cancer patients. Presently, much work is focused on the identification of useful biologic and molecular markers in the diagnosis and therapy of LSCC^7, 8^. In recent years, scholars have proposed that the combined expression of multiple genes may serve as a satisfactory marker for prognostic prediction. As tissue microarray technology and proteomics develop, more and more metastasis suppressor genes have been found. Exploring relations among anti-metastasis genes plays an important part in the prognostic prediction and treatment of LSCC. Therefore, it is imperative to find novel molecular markers underlying the tumorigenesis of LSCC for early diagnosis and prognostic assessment.

We undertook a retrospective analysis of data from a prospective longitudinal study of 170 patients over an extended time period (2002–2013) to examine the epidemiology of laryngeal squamous cell carcinoma (LSCC) with regard to age, laryngectomy, neck dissection, tumor differentiation, T-stage, N-stage, TNM stage and treatment modality. Meanwhile, in analyzing possible genetic polymorphisms associated with the susceptibility and prognostic evaluation of SLCC, we hoped to be able to identify possible points of intervention that may lead to improving patient survival.

## Materials and methods

### Subjects

In this study, we identified 170 patients without distant metastasis of LSCC who underwent partial or total laryngectomy. A total of 170 laryngeal cancer patients′ blood samples were randomly collected at the First Affiliated Hospital of Xi’an Jiaotong University School of Medicine between January 2002 and April 2013. The histologic diagnosis of tumors was made and agreed upon by at least two senior pathologists at the Department of Pathology of the Hospital based on World Health Organization (WHO) criteria. All of the patients are men and their ages ranged from 32 years to 82 years with an average of 60.75. All patients underwent a standard clinical examination and none of these patients had received any therapy before admission before surgery. All cases were systematically classified based on the Union of International Cancer Control (UICC, 2010) TNM staging system of laryngeal carcinomas, which essentially establishes the modality of therapy.

Following-up form including access to medical records and telephone contact. The medical records of these patients were reviewed to assess the patients’ characteristics, including age, laryngectomy (partial or total), neck dissection (yes or no), tumor differentiation, T-stage, N-stage, TNM stage and final status on the last follow-up examination. We chose April 2013 as the end time of eligibility with the goal of having adequate follow-up of individual participants with a median follow-up time of 48 months (range, 3 to 122 months).100 patients were identified as death and none of patients was lost from follow-up data. The survival time was defined as from the date of surgery to the date of death. The overall survival rate for five years in this group of patients was 47.1% (**Fig.1A**). Cases of death patients were regarded as censored data and marked on the survival curves. Informed consent was obtained from each patient according to protocols approved by the ethics committees of the First Affiliated Hospital of Xi’an Jiaotong University School of Medicine.

**Fig 1.**
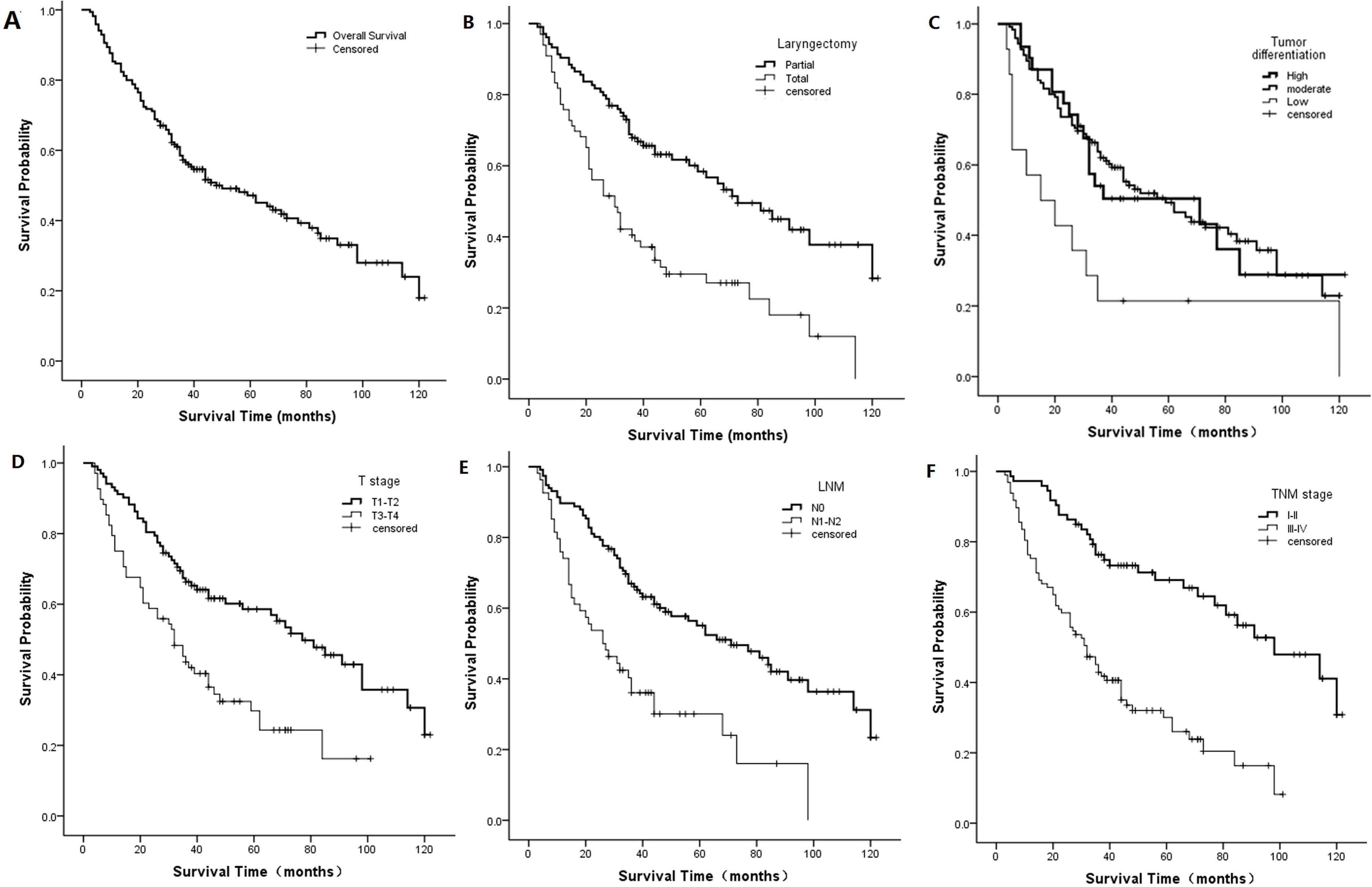
Kaplan-Meier curves of (A) overall survival; (B) laryngectomy (p = 0.000, log-rank test); (C) Tumor differentiation (p = 0.008, log-rank test); (D) T-stage (p = 0.000, log-rank test); (E) N-stage (p = 0.000, log-rank test); (F) TNM stage (p = 0.000, log-rank test). The graphs present 10 years of follow-up. The log-rank test was based on the full data.

### SNP selection and genotyping

We successfully selected 14 single nucleotide polymorphisms (SNPs) of 1,2 and 3 chromosome, which associated with cancer of the aerodigestive tract. Basic information of the 14 candidate SNPs in this study were shown in Table 1. We extracted genomic DNA from peripheral blood using a GoldMag-Mini Whole Blood Genomic DNA Purification Kit (GoldMag Ltd. Xi'an, China) depending on the manufacturer's protocol. Sequenom MassARRAY Assay Design 3.0 Software was used to design primers for amplification and extension reactions^9^. SNP genotypes using the standard protocol recommended by the manufacturer was performed by Sequenom MassARRAY RS1000^9^. Finally, Sequenom Typer 4.0 Software was used to perform the data management and analysis^9, 10^.

**Table 1.** Basic information of SNPs in this study

| SNP-ID | Position | Band | Allele | Gene | Role |
| --- | --- | --- | --- | --- | --- |
| rs17401966 | 10385471 | 1p36.22 | G/A | KIF1B | Intron |
| rs4072037 | 155162067 | 1q22 | G/A | MUC1 | Coding exon |
| rs1912453 | 162821291 | 1q23.3 | C/T | C1orf110 | Downstream |
| rs10911251 | 183081194 | 1q25.3 | A/C | LAMC1 | Intron |
| rs6687758 | 222164948 | 1q41 | G/A | -- | -- |
| rs11903757 | 192587204 | 2q32.3 | C/T | -- | -- |
| rs9288520 | 217481271 | 2q35 | A/G | -- | -- |
| rs966423 | 218310340 | 2q35 | T/C | DIRC3 | Intron |
| rs6736997 | 235615197 | 2q37.2 | A/C | -- | -- |
| rs975334 | 2846316 | 3p26.2 | C/T | CNTN4 | Intron |
| rs8180040 | 47388947 | 3p21.31 | A/T | KLHL18 | Downstream |
| rs9841504 | 114362764 | 3q13.31 | G/C | ZBTB20 | Intron |
| rs10936599 | 169492101 | 3q26.2 | T/C | ARPM1 | Promoter |
| rs2239612 | 186793242 | 3q27.3 | T/C | ST6GAL1 | Intron |

### Statistical analysis

Correlations between categoric variables were done using the chi-square test. Survival curves were drawn using the Kapla-Meier method. Differences between curves were analyzed using the log-rank test. A cox proportional hazards model was applied to estimate the risk by calculating hazard ratios (HR) and the 95% confidence intervals (CI) for categorical variables of exposure. Multivariate Cox regression analysis models were then developed that adjusted for the most important covariates, including SNP, Tumor differentiation, neck dissection, T stage, and TNM stage. A multivariate analysis taking into account the variables that were found to be significant on univariate analysis. Usually, prognostic impact factors with p ≤ 0.2 in univariate analysis were regarded as the candidate variables into the multivariate regression model. The SPSS statistical software package version 17.0 (SPSS Inc., Chicago, IL, USA) was used for all analyses. P values less than 0.05 were examined statistically significant.

## Results

### Patient characteristics and treatment outcomes

Briefly, from January 2002 and April 2013, a total of 170 LSCC patients in the First Affiliated Hospital of Xi’an Jiaotong University School of Medicine were recruited in this study. All the patients were males and the average age (mean ± SD) was 60.75 ± 10.082 years (range, 32 to 82 years). Patients were followed up since the diagnosis until the end of April, 2013. Among the 170 LSCC patients, none of patients were lost from follow-up data. Finally, 170 patients were investigated successfully. The median time of follow-up was 38 months (minimum and maximum were 3 and 122 months, respectively) and median survival time was 48 months. 100 out of them (58.82%) died of LSCC during the follow-up period. The overall survival rate for five years in this group of patients was 47.1% (**Fig.1A**).

As showed in Table 2, we had listed the possible factors that influence the prognosis. We further investigated the relationships of these possible factors stratification with prognostic survival by univariate analysis and survival curves were drawn using the Kapla-Meier method. We found that analyses stratified by patients’ laryngectomy (p = 0.000, **Fig. 1B**), differentiation (p = 0.008, **Fig. 1C**), T-stage (p = 0.000, **Fig. 1D**), N-stage (p = 0.000, **Fig. 1E**), TNM stage (p = 0.000, **Fig. 1F**) were associations with survival during the prognosis based on the log-rank test. However, prognostic survival was no correlations with age (p = 0.456) and neck dissection (p = 0.188) in stratified analyses. Compared with partial laryngectomy, high differentiation, T1-T2 stage, N0 stage and I-II TNM stage, the HR of total laryngectomy (95% CI, 1.576 - 3.492; p = 0.000), poor differentiation (95% CI, 1.150 - 4.997; p = 0.020), T3-T4 stage (95% CI,1.448 - 3.253; p = 0.000), N1-N2 stage (95% CI, 1.582 - 3.623; p = 0.000) and III-IV TNM stage (95% CI, 2.100 - 5.180; p = 0.000) increased to 2.346, 2.397, 2.170, 2.394, 3.298, respectively.

**Table 2.**
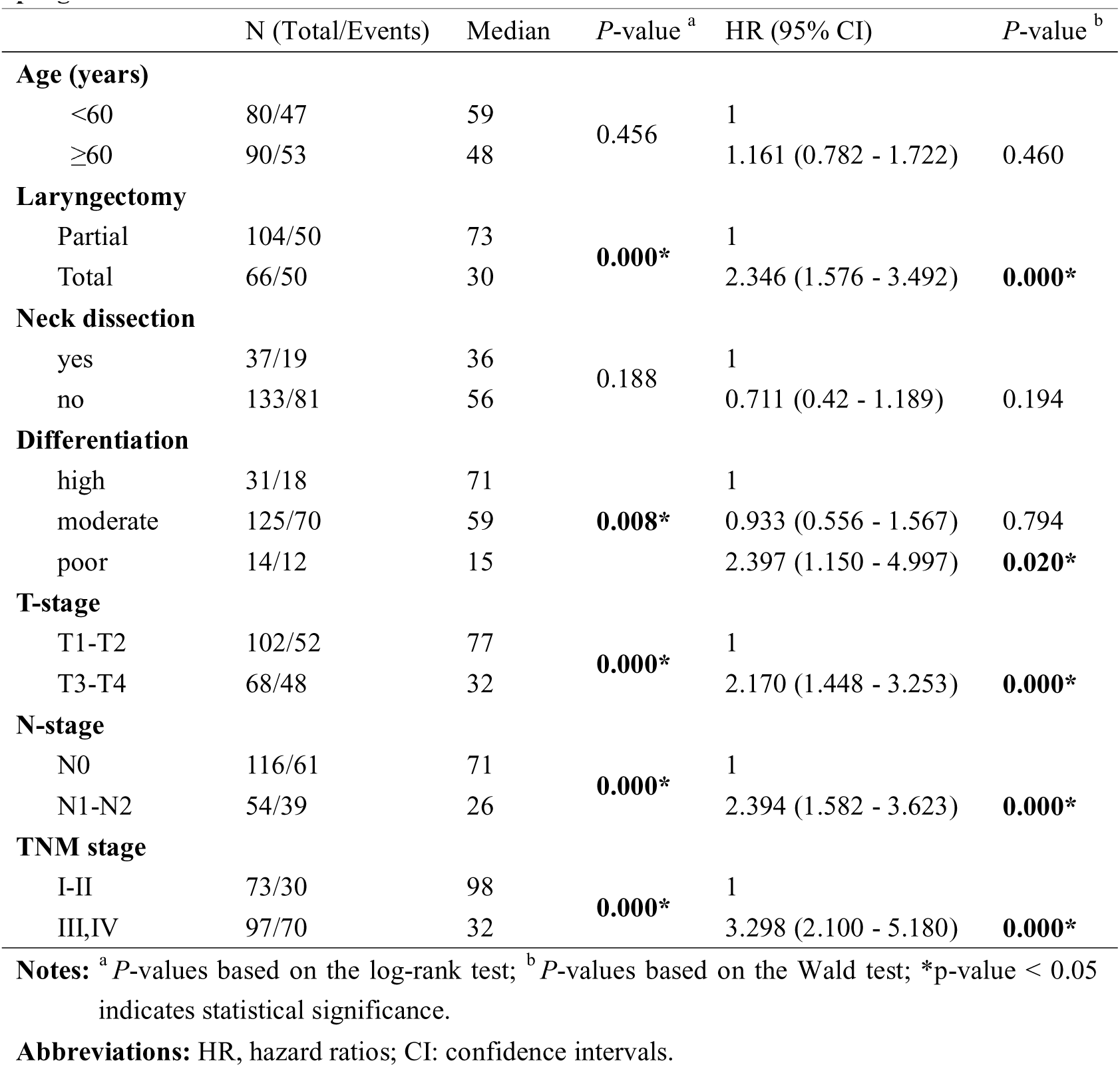
Patient characteristics and treatment outcomes - univariate associations with LSCC prognosis

### Genetic polymorphisms and outcome correlations

And then we assessed the associations between 14 SNPs genotypes and prognostic survival by univariate analysis, the results we found were listed in Table 3. Significant associations were detected between two SNPs (rs11903757, p = 0.021, **Fig. 2A**; rs966423, p = 0.000, **Fig. 2B**) genotypes and prognostic survival when the entire study patients were examined based on the log-rank test. For rs11903757 close to *NABP1*, the CT genotype (HR, 2.001; 95% CI, 1.091 - 3.673; p =0.025) resulted in a higher risk when compared to the T/T genotype. With regard to rs966423 in *DIRC3*, the prognostic survival was significantly worse in patients with TT genotype (HR, 7.721; 95% CI, 2.748 - 21.695; p =0.000) than those with the CC genotype, and the patients carried TC (HR, 1.089; 95% CI, 0 .701 - 1.690; p =0.705) genotype had no significant association with prognostic survival.

**Table 3.** Genetic polymorphisms and outcome - univariate associations with LSCC prognosis

| SNP-ID | genotype | N(Total/Events) | Median | *P*-value<sup>a</sup> | HR (95% CI) | *P*-value<sup>b</sup> |
| --- | --- | --- | --- | --- | --- | --- |
| rs17401966 | A/A | 125/73 | 48.0 |  | 1 |  |
|  | G/A | 36/19 | 66.0 | 0.437 | 0.905 (0.545 - 1.502) | 0.669 |
|  | G/G | 8/7 | 35.0 |  | 1.576 (0.724 - 3.430) | 0.252 |
| rs4072037 | A/A | 16394 | 46 |  | 1 |  |
|  | G/A | 2/2 | 81 | 0.843 | 0.976 (0.239 - 3.985) | 0.973 |
|  | G/G | 3/3 | 59 |  | 1.403 (0.442 - 4.450) | 0.565 |
| rs1912453 | T/T | 41/25 | 46 |  | 1 |  |
|  | C/T | 113/65 | 62 | 0.389 | 0.936 (0.589 - 1.486) | 0.779 |
|  | C/C | 15/10 | 35 |  | 1.489 (0.712 - 3.116) | 0.290 |
| rs10911251 | C/C | 45/28 | 36 |  | 1 |  |
|  | A/C | 81/48 | 59 | 0.720 | 0.831 (0.518 - 1.333) | 0.443 |
|  | A/A | 34/19 | 66 |  | 0.949 (0.530 - 1.700) | 0.860 |
| rs6687758 | A/A | 114/68 | 59 |  | 1 |  |
|  | G/A | 44/26 | 44 | 0.979 | 1.047 (0.664 - 1.651) | 0.842 |
|  | G/G | 5/2 | 34 |  | 0.975 (0.238 - 4.000) | 0.972 |
| rs11903757 | T/T | 156/88 | 59 |  | 1 |  |
|  | C/T | 14/12 | 23 | **0.021\*** | 2.001 (1.091 - 3.673) | **0.025\*** |
|  | C/C | -/- | - |  | - | - |
| rs9288520 | G/G | 92/52 | 48 |  | 1 |  |
|  | A/G | 72/45 | 50 | 0.786 | 1.085 (0.727 - 1.619) | 0.691 |
|  | A/A | 4/2 | 22 |  | 0.692 (0.166 - 2.877) | 0.613 |
| rs966423 | C/C | 111/65 | 62 |  | 1 |  |
|  | T/C | 52/29 | 44 | **0.000\*** | 1.089 (0.701 - 1.690) | 0.705 |
|  | T/T | 4/4 | 5 |  | 7.721(2.748 - 21.695 ) | **0.000\*** |
| rs6736997 | C/C | 105/66 | 44 |  | 1 |  |
|  | A/C | 64/33 | 62 | 0.540 | 0.878 (0.577 - 1.336) | 0.543 |
|  | A/A | -/- | - |  | - | - |
| rs975334 | T/T | 125/74 | 48 |  | 1 |  |
|  | C/T | 42/25 | 46 | 0.584 | 1.017 (1.017 - 1.603) | 0.941 |
|  | C/C | 3/1 | 59 |  | 0.373 (0.052 - 2.687) | 0.328 |
| rs8180040 | T/T | 51/32 | 44 |  | 1 |  |
|  | A/T | 51/28 | 62 | 0.754 | 0.836 (0.501 - 1.394) | 0.493 |
|  | A/A | 43/24 | 62 |  | 0.858 (0.505 - 1.458) | 0.572 |
| rs9841504 | G/G | 132/77 | 56 |  | 1 |  |
|  | G/C | 26/16 | 40 | 0.680 | 1.120 (0.652 - 1.925) | 0.682 |
|  | C/C | 3/2 | 26 |  | 1.763 (0.430 - 7.223) | 0.431 |
| rs10936599 | C/C | 48/34 | 44 |  | 1 |  |
|  | T/C | 73/37 | 62 | 0.153 | 0.635 (0.398 - 1.014) | 0.057 |
|  | C/C | 47/28 | 36 |  | 0.818 (0.494 - 1.356) | 0.436 |
| rs2239612 | C/C | 101/54 | 71 |  | 1 |  |
|  | T/C | 45/31 | 40 | 0.192 | 1.488 (0.954 - 2.321) | 0.080 |
|  | T/T | 12/8 | 44 |  | 1.332 (0.632 - 2.810) | 0.451 |

**Notes:** <sup>a</sup> *P*-values based on the log-rank test; <sup>b</sup> *P*-values based on the Wald test; \**p*-value < 0.05 indicates statistical significance.
**Abbreviations:** HR, hazard ratios; CI: confidence intervals

**Fig 2.**
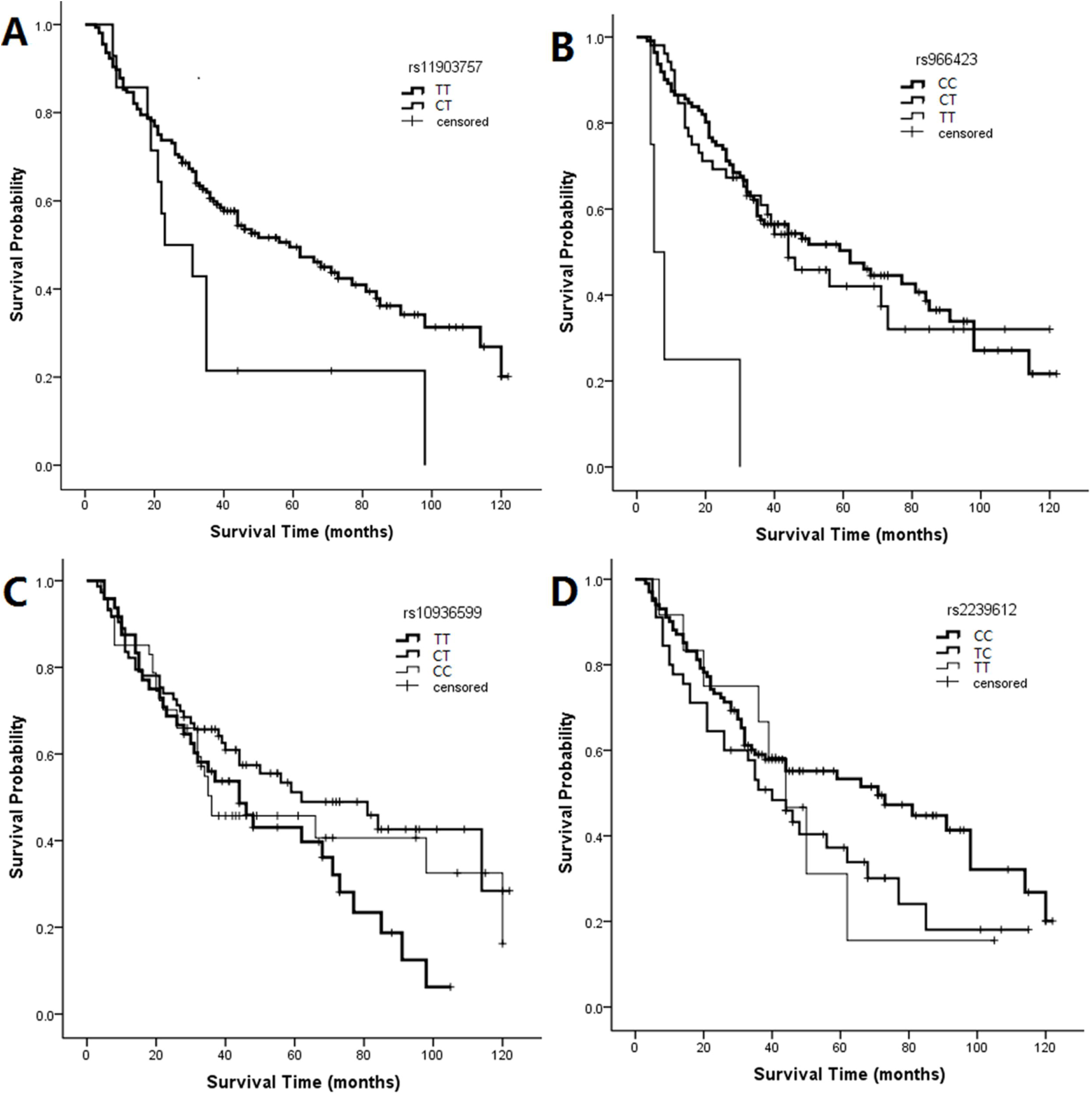
Kaplan-Meier curves of (A) rs11903757 genotypes (p = 0.021, log-rank test); (B) rs966423 genotypes (p = 0.000, log-rank test); (C) rs10936599 genotypes (p = 0.153, log-rank test); (D) rs2239612 genotypes (p = 0.192, log-rank test). The graphs present 10 years of follow-up. The log-rank test was based on the full data.

### Multivariate analysis of treatment outcome

Since the rs11903757 and rs966423 genotypes might be prognostic markers in SLCC, we performed univariate analysis first in SLCC patients. We found that patients with total laryngectomy, poor differentiation, T3-T4 stage, N1-N2 stage, III-IV TNM stage and SNPs had a worse prognostic survival by univariate analysis. Taking into account the rs10936599 (p = 0.153, **Fig. 2C**) in the promoter region of *ARPM1* and rs2239612 (p = 0.192, **Fig. 2D**) in the intron region of *ST6GAL1* might be associated with prognostic survival. We adjusted to the above confounding factors into a multivariate Cox proportional hazards model, calculating HR and 95% CI of different genotypes, which were evaluated to measure the impact of minimum allele on the prognosis of disease. We got similar results which were presented in Table 4. With regard to rs11903757, the genotype CT (HR = 2.036; 95% CI, 1.071 - 3.872; p = 0.030) increased HR more than 2-fold compared to genotype TT. For rs966423, the prognostic survival was significantly worse in patients with TT genotype (HR = 11.677; 95% CI, 3.901 - 34.950; p =0.000) than those with the CC genotype. Rs10936599 and rs2239612 were not related to the prognosis of disease.

**Table 4.** Genetic polymorphisms and outcome - multivariate associations with LSCC prognosis

| SNP-ID | Genotype | HR(95% CI) | *P*-value |
| --- | --- | --- | --- |
| rs11903757 | TT | 1 |  |
|  | CT | 2.036(1.071 - 3.872) | **0.030\*** |
| rs966423 | CC | 1 |  |
|  | TC | 0.154(0.727 - 1.833) | 0.543 |
|  | TT | 11.677(3.901 - 34.950) | **0.000\*** |
| rs10936599 | TT | 1 |  |
|  | CT | 0.780(0.475 - 1.280) | 0.326 |
|  | CC | 0.933(0.555 - 1.567) | 0.792 |
| rs2239612 | CC | 1 |  |
|  | TC | 1.232(0.779 - 1.949) | 0.373 |
|  | TT | 1.224(0.576 - 2.600) | 0.599 |

**Notes:** <sup>b</sup> *P*-values based on the Wald test; \**p*-value < 0.05 indicates statistical significance.
**Abbreviations:** HR, hazard ratios; CI: confidence intervals.

## Discussion

The second most common type of head and neck cancer is laryngeal cancer. New cases of laryngeal cancer diagnosed in the USA every year are estimated at 12,000. The incidence of laryngeal cancer is much higher in men than in women, especially for those found between 60 and 70 years old^6^. In current study, all of the patients are men. We found patients with total laryngectomy, poor differentiation, T3-T4 stage, N1-N2 stage, III-IV TNM stage had a worse prognostic survival and these findings are consistent with results of most other studies^11-14^. We also showed that the polymorphisms of rs11903757 near *NABP1* gene and rs966423 in *DIRC3* gene correlate with prognostic survival in patients with SLCC. We revealed that rs11903757 CT and rs966423 TT genotypes are unfavorable genotypes, and the presence of unfavorable allele resulted in a worse outcome in this patient group.

For rs11903757, an intergenic on chromosome 2q32.3, are in closest proximity to the *NABP1* gene (44 kb centromeric) and the gene serum deprivation response (112 kb telomeric), which encodes for the serum-deprivation response phosphatidylserine-binding protein^15^. The CT genotype was association with worse prognostic survival. The poor prognostic genotype unveiled in this study also predict inferior outcomes for other cancer type. Peters et al^16^ have recently reported a genome-wide significant association between rs11903757and colorectal cancer risk in a combined analysis of European and Asian case-control series. But another study^17^ reported that they failed to find convincing evidence for a previously published data with association between rs11903757 and colorectal cancer risk. The rs966423 is located in the intron region of the *DIRC3* at 2q35 and the TT genotype was associated with worse survival in SCLC patients. *DIRC3* (disrupted in renal cancer 3) was first identified in 2003. Its disruption in t(2;3)(q35;q21) translocation was observed in renal cell carcinoma^18^, and although the function of *DIRC3* is unknown, it is presumed to have tumor suppressor activity. In the previous GWAS, *DIRC3* was associated both with the risk of thyroid cancer and with the thyroid stimulating hormone level^19^. It is thus possible that changes in *DIRC3* alter thyroid stimulating hormone production and in an indirect way TC development as a result of decreased differentiation of the thyroid epithelium. Whether there was a similar mechanism in laryngeal cancer, we are not involved in the study. We will pay attention to these problems in the future research. Previous study reported that the presence of the rs966423 TT genotype was associated with a significant increase in overall mortality of patients with differentiated thyroid cancer^20^. The CT and CT + TT genotypes of rs966423 were more common in papillary thyroid cancer patients with extra thyroidal extension and more advanced T stage^21-23^. As we know, laryngeal carcinoma and thyroid carcinoma are the two main types of malignant tumors in the head and neck and both sides of the thyroid gland attached to the lower part of the throat. It has been reported that thyroid carcinomas were incidentally found in 0.7 percent to 3 percent of surgeries for another primary head and neck cancer of nonthyroid origin^24-29^. A clinically unexpected simultaneous thyroid cancer confirmed postoperatively from thyroid tissue partially removed with a laryngeal specimen for laryngeal cancer is rare^30^. Therefore, further study should expand sample size and involve in different regions or populations to study in a deepgoing way.

Of course, our research still has some limitations. First of all, there are only 170 patients in our trial for statistical analysis, the sample size is small, but also need a wide range of samples to verify our results. Secondly, our sample is limited to Shaanxi, China, and need to be validated in different regions and confirmed with larger sample size from different ethnic populations. In addition, it would be interesting to elucidate the functional relevance of the variants to get insight into the mechanism underlying the association.

In conclusion, this study provided new evidence for patients with total laryngectomy, poor differentiation, T3-T4 stage, N1-N2 stage, III-IV TNM stage had a worse prognostic survival and seemed to be the first that demonstrated rs11903757 GT genotype on chromosome 2q32.3 close to *NABP1* and rs966423 TT genotype in the intron region of *DIRC3* on chromosome 2q35 predict poor prognostic survival in patients with LSCC. It will have a guiding role in the future clinical treatment of laryngeal cancer and it may lead to improving patient survival. Further study would be interesting to elucidate the functional relevance of the variants to get insight into the mechanism underlying the association.

## Acknowledgements

This work is supported by China Postdoctoral Science Foundation (NO. 2015M572575) and Key Science and Technology Program of Shaanxi Province, China (N0.2014K11-01-01-09). We are grateful to all the patients and individuals for their participation.

## Abbreviations

LSCC: Laryngeal squamous cell carcinoma
SNPs: single nucleotide polymorphisms
HR: hazard ratios
CI: confidence interval

